# Exon-skipping and genetic compensation due to biallelic mutations in the neurodevelopmental disease gene *LNPK*

**DOI:** 10.1101/2025.05.30.656906

**Authors:** Rose M. Doss, Sara A. Wirth, Jonathan W. Pitsch, Caroline M. Dias, Andrea L. Gropman, Martin W. Breuss

## Abstract

Homozygous loss-of-function mutations in *LNPK*, the gene encoding the endoplasmic reticulum-associated protein lunapark, have previously been linked to an autosomal recessive neurodevelopmental syndrome. Here, we describe an individual harboring compound heterozygous predicted splice site mutations with an overall matching phenotype. In cultured fibroblasts, these mutations result in a dearth of transcript and severe loss of protein, thereby establishing their likely pathogenicity. The underlying reduction in gene expression is due to the activation of the nonsense-mediated decay (NMD) pathway as a consequence of exon-skipping rather than intron retention, leading to aberrant transcripts. We further demonstrate that *LNPK* is subject to genetic compensation, as both cells from the affected individual and her mother exhibit a significant increase in transcript compared to a control cell line when treated with an inhibitor of NMD. Together, this report describes novel disease-causing variants in *LNPK* and reveals their impact on transcription and mRNA stability.

## Introduction

The endoplasmic reticulum (ER) is a large and architecturally complex network of sheets and tubules that has specialized functions in protein synthesis and distribution, lipid synthesis, and calcium ion storage and release (1-3). The structural integrity of the ER is critical in maintaining cellular homeostasis and function, especially in neurons that rely on a more complex version of this network (4); consequently, genetic variants affecting ER morphology-related proteins result in an array of neurodevelopmental disorders (5-8).

*LNPK* encodes for lunapark, a conserved, membrane-associated, curvature-stabilizing protein within tubular three-way junctions of the ER (9-11). Underlining the role of ER structure in neurodevelopment, a recent CRISPR/Cas9-based screen evaluating the important contributors in brain development found that *LNPK* knockout resulted in significant deficits in saltatory movement in interneuron migration (12). Clinically, 20 neurodevelopmental disorder patients with homozygous loss-of-function mutations in *LNPK* have been previously identified; they presented with a range of atypical neurodevelopmental phenotypes including moderate to severe intellectual disability, epilepsy, movement disorders, global developmental delays, hypotonia, corpus callosum hypoplasia, and the structural change called “ear-of-the-lynx” sign (i.e., signal alterations of the forceps minor) on brain imaging (Table 1) (13-15).

**Table 1.**
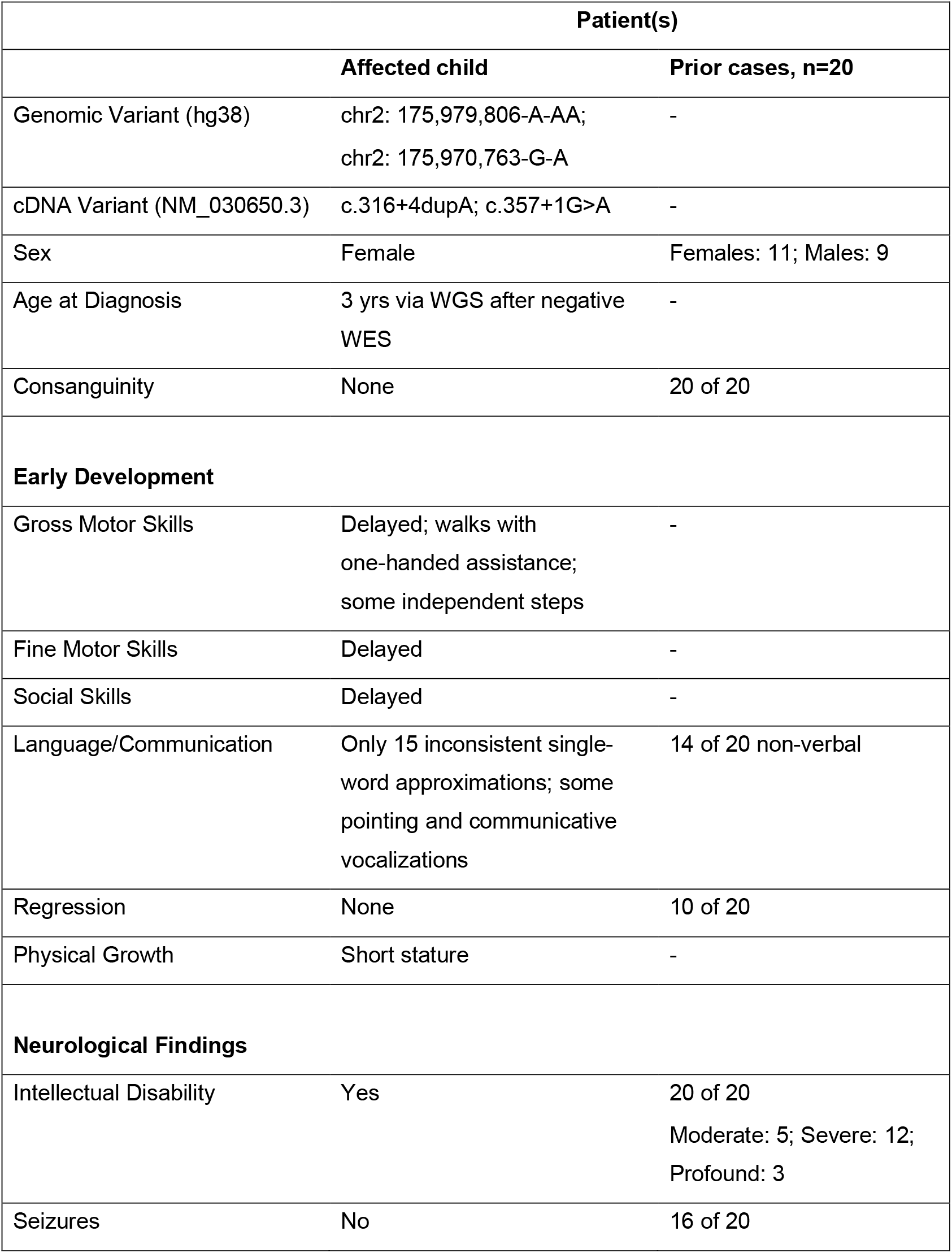
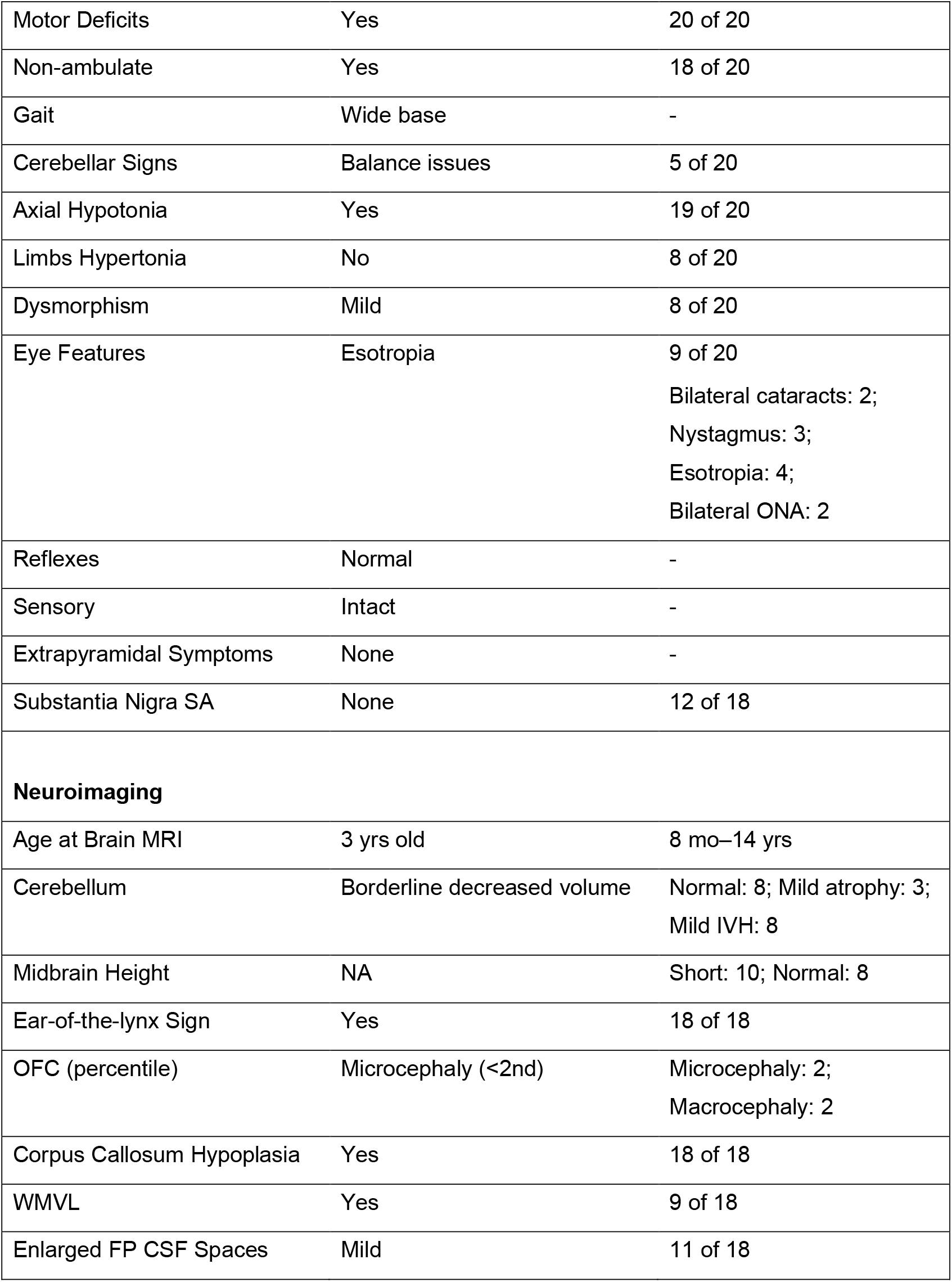
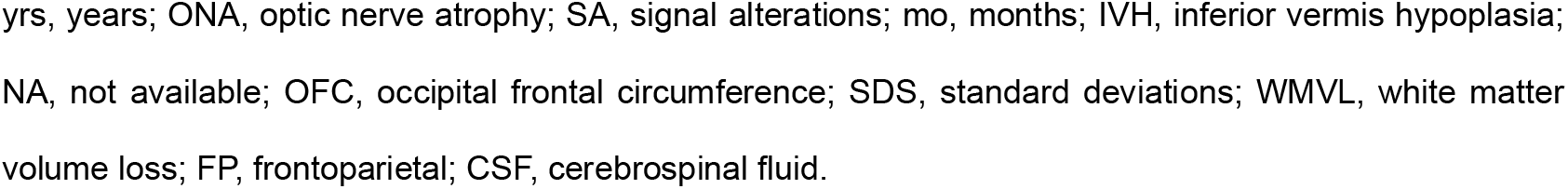
Clinical findings associated with *LNPK* variants.

Here, we report our findings from a child with biallelic variants in *LNPK* with a matching but potentially milder neurodevelopmental phenotype. The two detected variants are located in intronic splice site regions and were bioinformatically predicted to impact expression due to defective splicing. Our results reveal that these variants, indeed, cause exon skipping, which subsequently activates nonsense-mediated decay (NMD), leading to significantly reduced RNA and protein levels. Upon inhibition of the NMD pathway, we see a significant increase in *LNPK* RNA beyond typical levels, consistent with the phenomenon of genetic compensation.

## Materials, Subjects, and Methods

### Cell lines

Skin punch biopsies were obtained from the mother and child subjects with informed consent. Samples were consented under the IRB protocols 07-0386 (PIs Shaikh and Coughlin; CU Anschutz Medical Campus) and PRO6058 (PI Gropman). The employed control line was fibroblasts from a healthy human subject, WS1 (ATCC, Gaithersburg, MD, USA; Cat# CRL-1502, RRID: CVCL_2766). Control line cells were obtained from the Anschutz Medical Campus Cell Technologies Shared Resource (RRID: SCR_021982). Culture conditions are reported in the supplementary methods.

### Pharmacological inhibition of nonsense-mediated decay

Inhibition of NMD was performed by treating cells with 0.5 μM of hSMG-1 inhibitor 11j (MedChemExpress, Monmouth Junction, NJ, USA; Cat# HY-124719) in DMSO for 20 hours before harvest. Negative control was DMSO with no added drug. Samples were then washed 2x with PBS (1X) and collected for RNA extraction.

### Molecular species isolation

Gel extractions were performed using the Monarch DNA Gel Extraction Kit (New England Biolabs, Ipswich, MA, USA; Cat# T1020S), as per manufacturer’s protocol. Briefly, samples were prepared for PCR by mixing cDNA with primers targeting *LNPK*: forward on exon 4 and reverse on exon 7 (see Table S1). PCR cycles: 1x 95°C 1 min; 40x 95°C 15 sec, 58°C 15 sec, 72°C 15 sec; 1x 72°C 7 min; 105°C lid temperature. PCR products were run on a 2% gel, allowing for proper separation of bands before excision was performed; gel fragments were then subjected to the Monarch kit protocol. Results were analyzed with Sanger sequencing. TOPO cloning was performed using a TOPO TA cloning kit (Invitrogen, Carlsbad, CA, USA; Cat# 450030), as per manufacturer’s protocol. Briefly, we performed PCR reactions as described above, mixed PCR products with an open TOPO vector and transformed into the provided competent bacteria. Individual colonies were analyzed by isolating plasmid DNA, performing PCR targeting vector region around the insert, purification with ExoSap, and analysis with Sanger sequencing.

Additional experimental details are reported in the supplementary methods.

## Results and Discussion

Here we report our findings on a female child with novel compound heterozygous variants of unknown significance (VUS) within the intronic splice site regions of *LNPK* (NM_030650.3). At 20 months of age, the patient was evaluated by a neurogenetics team that performed chromosomal microarray and whole-exome sequencing; both tests failed to identify a pathogenic or likely pathogenic variant. Further testing using whole-genome sequencing (WGS) revealed compound heterozygous VUS in *LNPK*. These were a paternally inherited c.316+4dupA and a maternally inherited c.357+1G>A variant located at the 5’ splice site regions of intron 5 and 6, respectively (Fig. 1A–B). An initial analysis of these variants with SpliceAI (v1.3) returned scores of 0.80 donor loss for c.316+4dupA, and 0.98 donor loss for c.357+1G>A. These scores represent the probability that the variant affects splicing within a 500 bp region around it, with scores ≥0.8 representing variants with a high probability of having a splice-altering consequence (16). There were no other pathogenic or likely pathogenic variants reported from the WGS data. Compared to previously documented *LNPK* cases, the presentation of the individual included a similar phenotype as well as oral dysphagia, mild obstructive sleep apnea, and subtle dysmorphic features including a round face, close-set eyes, and epicanthal folds (see Clinical Report in supplementary materials). Of note, the patient has had no documented seizures or regression to date, indicating a possibly weaker deleterious effect of these variants than other reported cases. Structurally, brain imaging revealed mild corpus callosum hypogenesis and hypoplasia (Fig. 1C); the findings also included the stereotypical “ear-of-the-lynx” sign, found in all cases with the *LNPK*-related neurodevelopmental disorder tested so far (Fig. 1D).

**Fig. 1.**
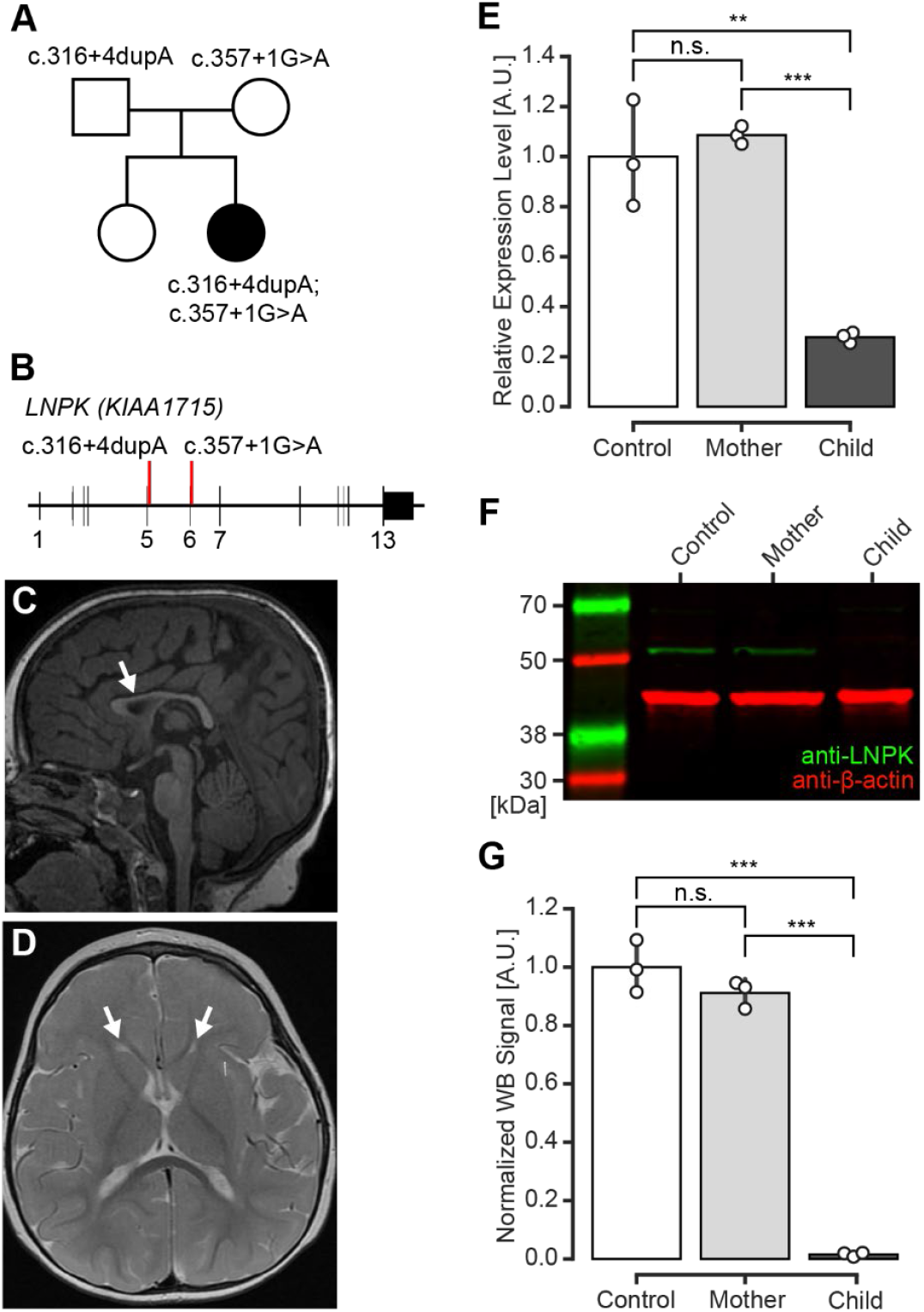
Phenotypic and molecular impact of two novel splice site mutations. **A** Family pedigree illustrating the genotype of each parent and the affected child. **B** Schematic of *LNPK* (NM_030650.3); red lines mark the location of the paternal variant (c.316+4dupA) within intron 5 and the maternal variant (c.357+1G>A) within intron 6. **C–D** The sagittal and axial MRI T2 weighted images of the affected child with arrows locating hypoplasia of the corpus callosum (C) and ear-of-the-lynx sign (D). Age at brain MRI was 3 years. **E** RT-qPCR expression levels were evaluated on control fibroblasts (control) and primary fibroblasts derived from the unaffected mother and the affected child. Expression is normalized to three housekeeping genes (see supplementary methods). **F** Western blot of protein lysates for each cell line. Expected molecular weights of lunapark and β-actin are 47.7 kDa and 42 kDa, respectively. **G** Quantification of each protein signal (Fig. S2). n=3 experimental replicates for (E) and (G). Barplots show the mean ± SEM and individual data points. n.s. p > 0.05, ** p < 0.01, *** p < 0.001 (pairwise comparisons with TukeyHSD confidence intervals).

To explore the potential impact and pathogenic mechanism of the two biallelic variants in more detail, we obtained primary fibroblasts from the unaffected mother and the affected child. We first confirmed the genotypes of the two resulting cell lines, as well as from a control WS1 cell line, at the *LNPK* loci by Sanger sequencing (Supplementary Fig. S1). Gene expression analysis via RT-qPCR and Western blot revealed that the mRNA (Fig. 1E) and protein levels (Fig. 1F–G) were significantly reduced in the child’s cell line, whereas the mother’s exhibited comparable levels of mRNA to the control. Thus, we conclude that the two biallelic splicing variants reduce *LNPK* expression to levels that are consistent with their pathogenicity.

We next hypothesized that these putative splice site mutations, consistent with the *in silico* prediction, could cause aberrant splicing and subsequent NMD, ultimately leading to the observed diminished protein product. To test this, we performed pharmacological inhibition of the NMD pathway using an SMG1-inhibitor (17). Upon this inhibition, we found a significant increase of mRNA levels in both the child and mother (Fig. 2A). To further explore the molecular mechanism causing NMD activation, we used NMD-inhibited samples—where aberrant transcripts persist— to resolve the transcripts by two approaches: 1. Targeted PCR on cDNA samples, followed by gel extraction and subsequent Sanger sequencing, or 2. Targeted PCR on cDNA samples followed by TOPO cloning, an additional round of targeted PCR, and subsequent Sanger sequencing. Together, these analyses revealed that each intronic variant resulted in single exon skipping (Fig. 2B–C), whereas the initially predicted intron retention was undetectable (Fig. S3). The exon-skipping events result in a frameshift, introducing a premature stop codon (see Data S1), which in turn likely triggers the activation of NMD. The potentially milder phenotypic presentation (i.e., the absence of epilepsy and regression) could be due to incomplete penetrance of the splicing defect. While the protein levels in fibroblasts appear almost completely diminished, which is inconsistent with this prediction, we were unable to assess tissue-specific expressivity and sensitivity to the impact of these splice site mutations (18).

**Fig. 2.**
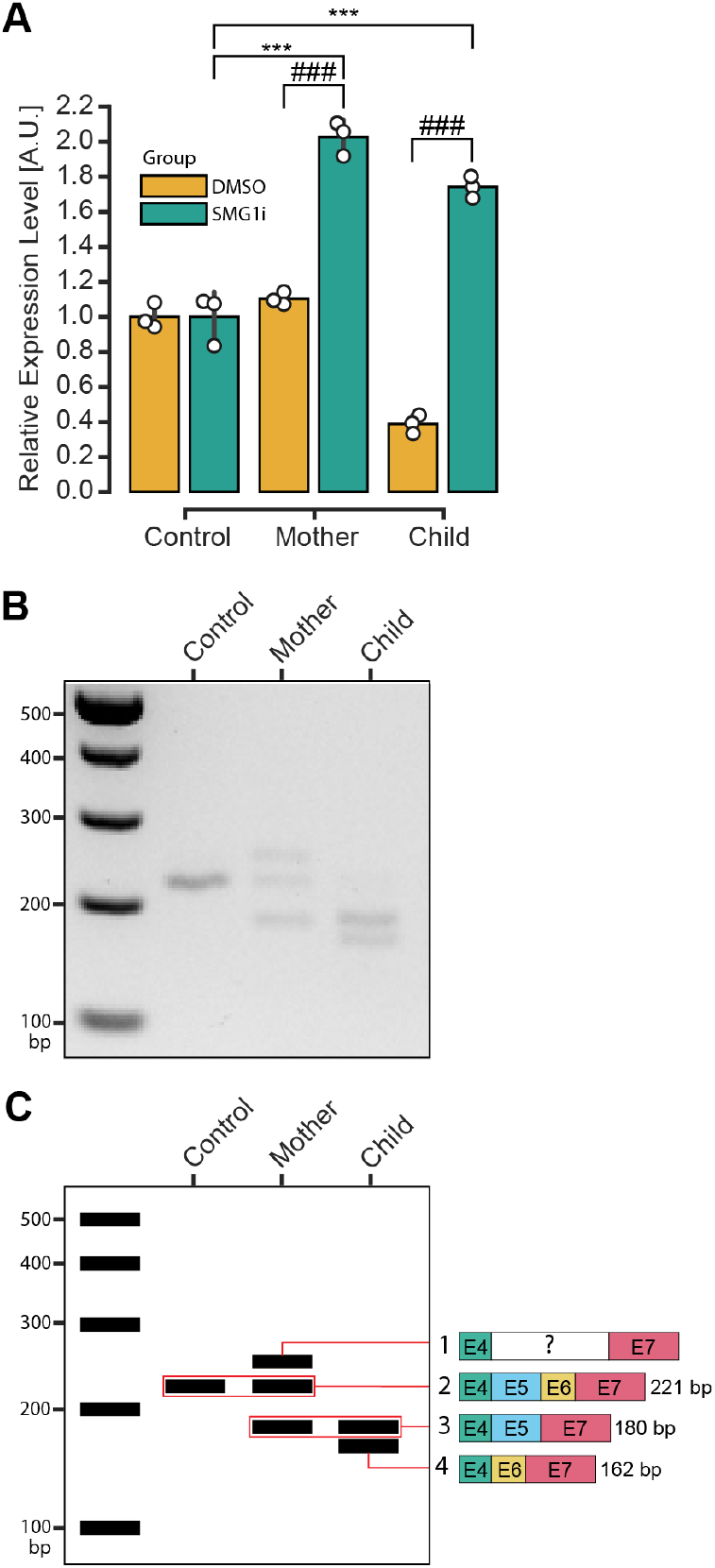
Impact of variants on mRNA stability and splicing. **A** RT-qPCR expression levels after each fibroblast cell line was exposed for 20 hours to either DMSO only (orange) or an SMG1-inhibitor (SMG1i, green) at 0.5 μM in DMSO. Expression is normalized to three housekeeping genes (see supplementary methods). Barplot shows the mean +/-SEM and individual data points (n=3 experimental replicates). *** or ### p < 0.001 (two-way ANOVA and pairwise comparisons with TukeyHSD confidence intervals). **B** RT-PCR on NMD-inhibited samples. **C** Visual summary of direct sequencing results from (B). Exon 7 in the mother’s top band was only partially resolved in the sequencing data. Sequences can be found in supplementary files (Data S1).

Surprisingly, the mRNA levels in both the mother and affected child significantly surpass the control cell line’s upon NMD inhibition, while the mother’s mRNA and protein levels were at a similar level to the control before treatment. This finding suggests a genetic compensation mechanism at play, where the upregulation of the functional gene copy in the mother restores protein levels, allowing for the normal physiological function of lunapark (19, 20). There are no known paralogs for *LNPK* in humans, therefore, the upregulation of mRNA is expected to come exclusively from both copies of *LNPK*. Additionally, despite several attempts using both molecular species isolation approaches, the mother’s top band (Fig. 2C) consistently could not be resolved, a result that may be due to non-specific primer binding due to the combined presence of fragments 2 and 3.

This is the first case report studying the molecular mechanisms of compound heterozygous variants in *LNPK* located near the donor splice sites for introns 5 and 6. Three previous cases had homozygous splice site variants reported, although the molecular impact was not explored. These findings could guide potential therapeutic approaches in this or related cases, suggesting that molecular assessment of predicted splicing defects remains an important addition to validation workflows. Overall, our study supports that the molecular impact of the two novel *LNPK* variants c.316+4dupA and c.357+1G>A is consistent with their pathogenicity. The potentially milder phenotype presentation compared to the previously identified cases is possibly due to an incomplete loss of protein function.

## Supporting information

Supplementary Materials (Figures, Table, Clinical Report, Methods)

Data S1 FASTA of nucleotide sequences from RT-PCR and predicted variant transcripts

## Data Availability Statement

All relevant data is provided in the supplementary files.

## Code Availability Statement

There is no relevant code generated for this study.

## Acknowledgments

We would like to thank the family who contributed samples and records to this study, Ana Mencia Moreno Chaza for assistance in preparing the clinical table and imaging figure included in this manuscript, and Dr. Sujatha Jagannathan for helpful discussions and guidance for the NMD-inhibition experiments. M.W.B. and C.M.D. acknowledge the Boettcher Foundation for their financial support.

## Author Contribution Statement

R.M.D. planned and executed experimental work with the support of S.A.W. R.M.D. analyzed and interpreted genetic results and data with support of J.W.P. A.L.G. and C.M.D. performed interpretation of clinical results. A.L.G. acquired biological samples. M.W.B. conceived and supervised the study. R.M.D. and M.W.B. wrote the manuscript and all authors saw and commented on it.

## Ethical Approval

IRB protocol 07-0386 (PIs Shaikh and Coughlin; CU Anschutz Medical Campus) and IRB protocol PRO6058 (PI Gropman).

## Supplementary Files

Supplementary Materials: Figures S1–S3, and Table S1 Supplementary Text: Clinical Report, and Supplementary Methods

Data S1 FASTA of nucleotide sequences from RT-PCR and predicted variant transcripts

