## Supplementary Materials (Figures, Table, Clinical Report, Methods) for "Exon-skipping and genetic compensation due to biallelic mutations in the neurodevelopmental disease gene *LNPK*"

Supplementary Figures

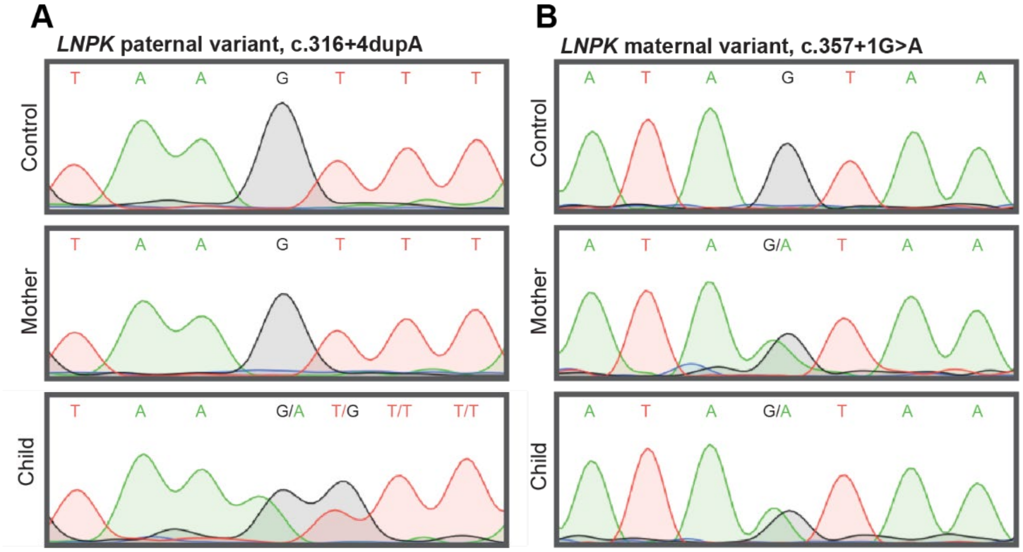


**Fig. S1 Confirmation of genotypes with Sanger sequencing.** Chromatogram showing the targeted variant regions within introns 5 and 6 of the *LNPK* gene for each sample. Sequencing results verify that the paternal variant, c.316+4dupA, is only present in the child while the maternal variant, c.357+1G>A, is present in both the mother and child.

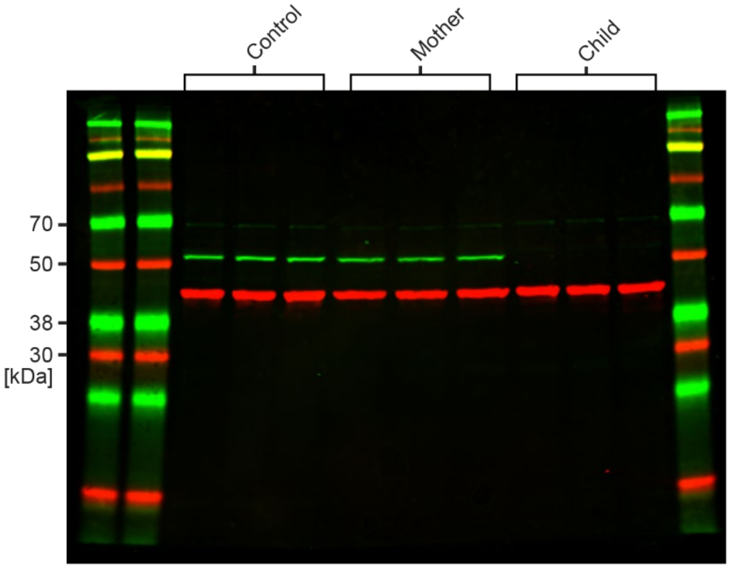


**Fig. S2 Western blot assay.** Western blot assessment illustrates a significant decrease of lunapark protein in the affected child. Expected molecular weights of lunapark and β-actin are 47.7 kDa and 42 kDa, respectively. n=3 for each sample. Blot underlies the quantification in Figure 1G.

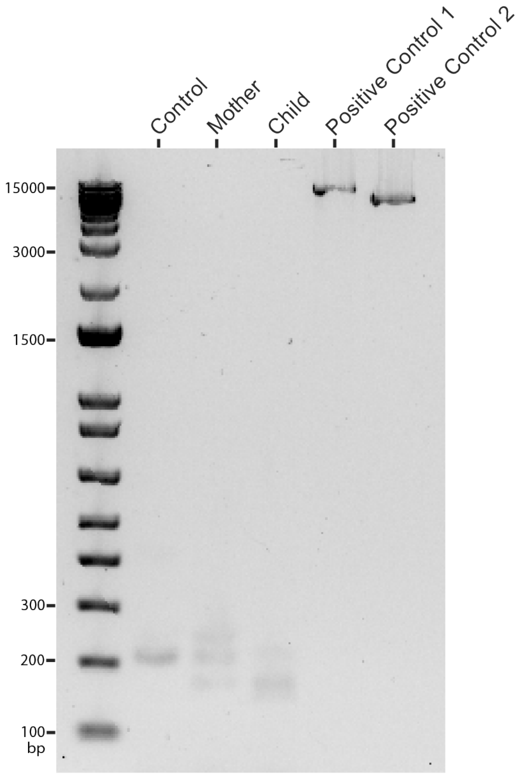


**Fig. S3 Intron retention analysis using long-range PCR.** Gel imaging of long-range PCR tests on each sample did not detect any evidence of intron retention. Two positive controls were performed on gDNA with primers designed to capture intronic and exonic regions around the target variants. Positive control 1 product = 9,649 bp. Positive control 2 product = 7,086 bp.

**Supplementary Table S1 Primer sequences used in this study**

| | |
|---|---|
| LNPK_SNV1_F | TCCCTTTTAGCATCTGGAGCA |
| LNPK_SNV1_R | GACAGAGTCAGGGGCATAGG |
| LNPK_SNV2_F | GTGTTTGCTACCGTGTGTCA |
| LNPK_SNV2_R | AAATGAGTCATATTGGAGGCCT |
| LNPK_exon4_F | ACTTGCCATGACACTCCCAT |
| LNPK_exon7_R | TCCGGACTCAAAGAAAGCAA |
| LNPK_intron5_R | TGTGCTGGTAAGACAGGTTG |
| LNPK_intron6_R | ACTCTCCAAGGAAGAAATATGCT |
| LNPK_intron6_F | CCATTATTGAAGTTGGTGCATGT |
| LNPK_intron5_F | GTGTTTGCTACCGTGTGTCA |
| LNPK_5'_qPCR_F | TGGGTGGATTATTTTCTCGATGG |
| LNPK_5'_qPCR_R | ACTTGCCATGACACTCCCAT |
| LNPK_3'_qPCR_F | TGCTTTCATCAGACAACCAG |
| LNPK_3'_qPCR_R | GATAATACAGAGCAGACAGATGAC |
| LNPK_LR_Intron-Exon5_F | CCCTTTTAGCATCTGGAGCA |
| LNPK_LR_Intron6_R | GTGGGGCTAGTTGATGCATC |
| LNPK_LR_Intron5 _F | GCCAACCTATATAAGCGGGC |
| LNPK_LR_Intron8 _R | GTGTGTTGGAGGTGTGCAAA |
| hsHPRT_qPCR_F | TGACACTGGCAAAACAATGCA |
| hsHPRT_qPCR_R | AGCTTGCTGGTGAAAAGGACC |
| hsPGK1_qPCR_F | ATGGATGAGGTGGTGAAAGC |
| hsPGK1_qPCR_R | AGTCAGCCATGTGAGCACTG |
| hsTBP_qPCR_F | TGCACAGGAGCCAAGAGTGAA |
| hsTBP_qPCR_R | TGGTGGGGAGCTGTGATGTG |

### Supplementary Text

Clinical Report

The proband was born to a 31-year-old female at 41 weeks after an uncomplicated pregnancy and delivery. Mother had one previous pregnancy that was also carried to a viable gestational age, delivered healthy, and has no relevant phenotypes to date. During pregnancy, the mother did not have any illnesses, and there was no exposure. Prenatal screens were negative, and fetal movement was reported as normal. The child was large for gestational age despite the absence of maternal gestational diabetes mellitus. There were no complications requiring neonatal intensive care unit admission, and there were no traumatic or ischemic events at birth. Apgar scores were 8 and 9. Following a two-day stay in the hospital, the mother and child were discharged home. Although the mother encountered postpartum hemorrhage, both she and the baby remained healthy. Furthermore, comprehensive screenings for neurological, musculoskeletal, hearing, and critical congenital heart disease were all within normal limits. The patient’s weight at birth was 4.31 kilograms (98$^{th}$ percentile), and the length at birth was 50.8 centimeters (81$^{st}$ percentile).

The patient’s family history revealed one 4-year-old sister who was developmentally typical and no family history of autism, hearing disability, or epilepsy. However, the maternal uncle had a history of learning disability and dyslexia. There was a family history of Duchenne’s muscular dystrophy (DMD) of unknown genetic etiology, affecting paternal cousins who had died due to the condition. Paternal great-aunts were suspected carriers of DMD.

The child presented with developmental delay as a primary concern, with motor and speech delays. At 6 months of age, she had begun rolling but could not walk at 20 months of age. She had only a few single words at 18 months of age and 8–10 consonant sounds. At 20 months of age, the patient was first seen by the neurogenetics team. She exhibited delays in gross motor skills, receptive and expressive language, along with hypotonia and episodes of choking. Mild brisk hyperreflexia and subtle dysmorphic features raised concerns regarding a potential central neurologic condition.

She was diagnosed with strabismus in an ophthalmological assessment at 16 months old. A later ophthalmological assessment at 2.5 years old revealed moderate esotropia and bilateral hyperopia, which was corrected with glasses. Additionally, the patient had a history of choking episodes with solids at least twice weekly. At the initial neurology visit, no concerns were noted in genitourinary, endocrine, hematological, or dermatological evaluations. Musculoskeletal assessment revealed gross motor delay and low tone, while neurological examination indicated hypotonia, and muscle weakness. There was no history of seizures. At that time, she was receiving physical therapy (PT), occupational therapy (OT), and speech therapy.

At 23 months of age, she underwent a modified barium swallow showing oral dysphagia characterized by reduced bolus formation with thin liquids and regular consistency. She had one episode of flash penetration with large sips of thin liquids but no aspiration.

Biochemical lab results were obtained during the initial neurology visit at 20 months old. These included plasma amino acids, urine organic acid, carnitine, acylcarnitine, thyroid-stimulating hormone, creatine phosphokinase, and lactate level. All were within normal range or non-diagnostic.

Subsequent brain imaging was obtained. First MRI at age 3 revealed generalized mild myelination delay versus hypomyelination. Additionally, the findings included a mildly simplified cerebral gyral and sulcal pattern, mild corpus callosum hypogenesis and hypoplasia, mild cerebral white matter and brainstem volume loss versus hypoplasia, and borderline decreased volume of the cerebellum. Moreover, mildly hypoplastic olfactory bulb and tracts were noted.

A follow-up MRI was obtained within a year to evaluate myelination status. The imaging revealed myelination had progressed in the interim from prior MRI but remained mildly deficient for age. Findings also suggested superimposed mild gliosis involving frontal and parietal white matter.

An EEG was obtained due to staring spells at 3.5 years of age. The EEG background displayed slow activity alongside a dysmature sleep pattern but no epileptiform activity. Genetic testing, including chromosomal microarray (CMA) and whole-exome sequencing (WES), yielded negative results. However, further testing using whole-genome sequencing (WGS) identified two heterozygous variants of uncertain significance in the *LNPK* (LUNAPARK) gene (NM_030650.3): one maternally inherited (c.357+1G>A) and one paternally inherited (c.316+4dupA). A skin biopsy was performed on the patient (at 3 years old) and the mother for research purposes.

At four years of age, the patient presented with continued hypotonia, oral motor dysfunction, mild dysmorphic features including a round face, close-set eyes, and epicanthal folds. There was no report of developmental regression. She ambulated with a posterior walker. She had ongoing delays in speech and communication. She could articulate five single words, replicate animal sounds and had delays in functional communication. When expressing her needs, she resorted to vocalizations, though she had one to two signs.

Most recent clinical evaluation was performed at the age of five. Physical exam identified short stature ($3^{rd}$ percentile for height; Z = -1.88; based on CDC, Girls, 2–20 Years), and reduced head circumference (below $2^{nd}$ percentile; Z <-2.05; based on Nellhaus, Girls, 2–18). Upon neuropsychological evaluation, the patient continues to make developmental gains. However, the pace of her development is slower than expected for her age. Additionally, caregiver ratings of her adaptive functioning and her independence skills are below age expectations. Therefore, the patient's cognitive profile is best described by the DSM-5 diagnosis of Intellectual Disability.

The patient experiences nocturnal disturbances, including nocturnal awakenings, snoring, and increased daytime sleepiness. A sleep study was performed and identified mild obstructive sleep apnea. Furthermore, there was no distinct REM sleep identified; however, this may have been underestimated due to high levels of beta frequency activity present on the EEG portion of the study. The patient was referred for actigraphy for further evaluation and treatment of sleep disturbances after a melatonin trial yielded no improvements in nocturnal awakenings.

Currently, the following steps for phenotype management are taken: The patient undergoes speech therapy, occupational therapy, and physical therapy. The patient's feeding habits are managed cautiously by her parents due to mouth muscle tone issues, with a preference for small food pieces given incrementally to prevent engorgement and choking. She receives feeding therapy for this issue and pica. The parents report no major behavioral issues currently, although she has some history of hair pulling and biting (self-injurious and preferred adults in close proximity). The parents report that the patient has a very positive affect and is easily excitable.

### Supplementary Methods

#### *Whole-Genome Sequencing (WGS)*

WGS was performed by Variantyx on family trio saliva samples from father, mother, and the affected child. Sequencing analysis revealed heterozygous paternally inherited c.316+4dupA, and heterozygous maternally inherited c.357+1G>A variants in the child's *LNPK* (NM_030650.3). These variants were classified as variants of uncertain significance (VUS). There were no other prioritized variants identified in other candidates or in the American College of Medical Genetics and Genomics (ACMG) list of genes to be reported as secondary findings that could impact medical management and decision making.

*Cell culture*

Primary fibroblast lines were established from skin punch biopsies (1). In short, biopsy was washed in triplicate with PBS (1X) (Thermo Fisher Scientific, Waltham, MA, USA; Cat# 10-010-023), cut into small pieces, and given sufficient time to adhere to tissue culture-treated flasks (Thermo Fisher Scientific, Cat# FB012935) before primary culture media was added. All fibroblast lines were grown at 37°C and 5% $CO_2$ in incubator in Minimal Essential Media (Thermo Fisher Scientific; Cat# 11095080) with 20% fetal bovine serum (Thermo Fisher Scientific; Cat# FB12999102), penicillin-streptomycin (1X) (Genesee Scientific, El Cajon, CA, USA; Cat# 25-512), and Amphotericin B (0.1X) (HyClone Amphotericin B (Fungiezone); Cytiva, Marlborough, MA, USA; Cat# SV30078.01). During the establishment of primary fibroblast lines from biopsies, mycoplasma removal agent (1X) (AbD Serotec Mycoplasma Removal Agent; Thermo Fisher Scientific; Cat# NC9369822) was added to the media. Cells were passaged when they reached 80–100% confluency. Passages were performed by removing media, washing twice with PBS (1X) and adding 0.25% trypsin-EDTA (1X) (Thermo Fisher Scientific; Cat# 25200056). Technical replicates were used for all experiments by growing each cell line in triplicate in 6-well plates (USA Scientific, Ocala, FL, USA; Cat# CC7682-7506). All experiments were performed on cells that were passaged fewer than 10 times.

*DNA extraction and genotyping*

DNA was obtained from pelleted cells of each fibroblast cell line using the DNeasy Blood & Tissue Kit (QIAGEN, Valencia, CA, USA; Cat# 69506), as per manufacturer's protocol. In short, cells that were more than 80% confluent were collected for DNA extraction. PCR was performed using GoTaq G2 Colorless Master Mix (Promega, Madison, WI, USA; Cat# M7833). Primers were targeted around variants of interest and resulting PCR products were purified with ExoSap (Exonuclease I; New England Biolabs, Ipswich, MA, USA; Cat# M0293L; Shrimp Alkaline Phosphatase (rSAP); New England Biolabs; Cat# M0371S) and sent for Sanger sequencing. Primer sequences used are found in Table S1.

*RNA extraction, cDNA synthesis, and RT-qPCR*

RNA extraction and purification on each cell line was performed with the RNeasy Mini Kit (QIAGEN; Cat# 74104) according to the manufacturer's handbook. Briefly, cells that were more than 80% confluent were collected for RNA extraction. After RNA purification, samples were standardized to the same approximate concentrations and subsequently underwent cDNA synthesis using the SuperScript III First-Strand synthesis system for RT-PCR (Invitrogen, Carlsbad, CA, USA; Cat# 18080-051) using the Oligo(dT) priming method and stored at -20°C until further analysis. RT-qPCR was performed using PowerTrack SYBR Green Master Mix for qPCR (Fisher Scientific, Waltham, MA, USA; Cat# A46113) on a Bio-Rad cycler, in conjunction with negative controls. Internal controls were housekeeping genes *HPRT*, *TBP*, and *PGK1*, and all reactions were performed in triplicate (2). To determine the relative mRNA levels, the geometric mean of the control genes was determined, and the delta CT was calculated (3). Primers used can be found in Table S1.

*Long-range PCR reactions*

Long-range PCR was performed using LongAmp Hot Start *Taq* 2X Master Mix (New England Biolabs, Ipswich, MA, USA; Cat# M0533S). Cycles: 1x 94°C 1 min, 30x 94°C 30 sec, 62°C 30 sec, 65°C 8 min, 1x 65°C 10 min; 105°C lid temperature. Primers used can be found in Table S1.

*Western blotting*

Protein extractions were obtained from technical replicates of each cell line and analyzed by Western blot using standard protocols. Briefly, this involved washing cells 1x in cold PBS (1X), and addition of cold RIPA Lysis and Extraction Buffer (Thermo Fisher Scientific, Waltham, MA, USA; Cat# 89900) with added protease inhibitor (Thermo Fisher Scientific; Cat# A32953). Samples were vortexed and 1 volume of Sample Buffer, Laemmli (2X) (Sigma-Aldrich, Carlsbad, CA, USA; Cat# S3401-10VL) was added, followed by heating at 95°C for 1 min before storage at -20°C until further analysis. Antibodies used were anti-LNPK (Sigma-Aldrich; Cat# HPA014205, RRID:AB_2234186, diluted to 0.2 μg concentration) and anti-beta Actin (Santa Cruz Biotechnology, Dallas, TX, USA; Cat# sc-47778, RRID:AB_626632, diluted to 0.13 μg). Quantification of protein signal intensity for each sample was determined with ImageJ/Fiji software (v1.54g). In short, the western blot image was split into red and green channels, inverted, and each band was isolated within the same rectangular area. This area was converted into peak signals, and the area under the peak was determined after removal of background signal. Lunapark signals were normalized to the β-actin loading control.

*Sanger sequencing*

Sanger sequencing was performed by Quintara Biosciences. Sequence data was visualized using SnapGene Viewer (SnapGene v7.2.1).

*Data analysis*

Experimental data was analyzed within Visual Studio Code (v1.94.2) using Python (v3.12.3). Data visualization and statistics were performed with the following packages: seaborn (v0.12.2) (4), pandas (v2.2.1) (5), matplotlib (v3.8.4) (6), numpy (v1.26.4) (7), scipy (v1.13.0) (8), and statsmodels (v0.14.0) (9).
